# Under the shadow: Old-biased genes are subject to weak purifying selection at both the tissue and cell type-specific levels

**DOI:** 10.1101/2022.09.21.508695

**Authors:** Nisan Yıldız, Hamit İzgi, Firuza Rahimova, Umut Berkay Altıntaş, Zeliha Gözde Turan, Mehmet Somel

## Abstract

The mutation accumulation theory predicts that aging is caused by accumulation of late-acting deleterious variants in the germ-line, due to weak purifying selection at old age. In accordance with this model, we and others have shown that sequence conservation among old-biased genes (with higher expression in old versus young adults) is weaker than among young-biased genes across a number of mammalian and insect species. However, questions remained regarding the source and generality of this observation. It was especially unclear whether the observed patterns were driven by tissue and cell type composition shifts or by cell-autonomous expression changes during aging. How wide this trend would extend to non-mammalian metazoan aging was also uncertain. Here we analyzed bulk tissue as well as cell type-specific RNA sequencing data from diverse animal taxa across six different datasets from five species. We show that the previously reported age-related decrease in transcriptome conservation (ADICT) is commonly found in aging tissues of non-mammalian species, including non-mammalian vertebrates (chicken brain, killifish liver and skin) and invertebrates (fruit fly brain). Analyzing cell type-specific transcriptomes of adult mice, we further detect the same ADICT trend at the single cell type level. Old-biased genes are less conserved across the majority of cell types analyzed in the lung, brain, liver, muscle, kidney, and skin, and these include both tissue-specific cell types, and also ubiquitous immune cell types. Overall, our results support the notion that aging in metazoan tissues may be at least partly shaped by the mutation accumulation process.

## Introduction

Aging is defined as progressive decline in fitness with increasing age (Kirkwood & Austad, 2000; López-Otín *et al*., 2013). The mutation accumulation theory provides a simple and powerful explanation of how aging could evolve (Medawar, 1952). This is based on the premise that in many natural populations, the number of individuals contributing to the next generation declines with age by extrinsic mortality, even without invoking the intrinsic effects of aging. Consequently, phenotypically deleterious variants that show their deleterious effects only at old ages, when the residual effective population size is low, can drift to fixation, creating the so-called “selection shadow” at late ages [reviewed in (Flatt & Partridge, 2018)]. The accumulation of such late-acting deleterious variants could then explain the prevalence of aging phenotypes, leading to Gompertzian mortality curves at old ages. Conversely, aging could be absent in species in which the probability of extrinsic mortality decreases with age, e.g. in species where individuals can constantly grow in size. This prediction is supported by empirical evidence: in diverse animal species, mortality can be stable or even decline through lifetime (Jones *et al*., 2014; Cohen, 2018).

Testing the role of the mutation accumulation mechanism on aging initially relied on studying inter-individual heterogeneity of age-related fitness reduction, a prediction of the model (Charlesworth & Hughes, 1996; Shaw *et al*., 1999; Wilson *et al*., 2007; Escobar *et al*., 2008). More recently, researchers have also begun employing molecular data to test the idea. For instance, Rodriguez and colleagues used genetic disease and polymorphism data in humans to show that late-acting disease variants segregate at higher frequencies than early-expressed variants (Rodríguez *et al*., 2017).

Yet another line of studies has combined sequence divergence data (*dN/dS* ratios; ratio of rate of non-synonymous mutations to synonymous mutations) and transcriptome data, under the assumption that late-expressed genes will carry late-acting variants. These studies showed that genes with increased expression levels in old versus young adults (old-biased genes) tend to be less conserved than young-biased genes in human brain aging (Somel *et al*., 2010), and across human bulk tissues (Jia *et al*., 2018). Recently, we systematically investigated the pattern of lower evolutionary conservation of genes expressed at late age using 66 transcriptome datasets representing human, macaque, mouse and rat bulk tissue samples (Turan *et al*., 2019). We identified the same trend, which we termed age-related decrease in transcriptome conservation (ADICT), in 76% of these datasets. Interestingly, although some tissues, such as brain, lung and liver, showed conspicuous ADICT signatures, other tissues, such as muscle and heart, revealed no consistent pattern. We further found old-biased and weakly conserved mammalian genes to be enriched in apoptosis and inflammation-related processes.

More recently, Cheng and Kirkpatrick reported low conservation of old-biased genes across human, mouse, fruit fly and mosquito transcriptomes (Cheng & Kirkpatrick, 2021), also showing that old-biased genes carry higher levels of functional polymorphism (pN/pS) and tend to be evolutionarily younger. Meanwhile, Harrison and colleagues studied sequence conservation of young- and old-biased genes identified in whole-body transcriptomes in relatively long-lived ant queens (*Cardiocondyla obscurior*) (Harrison *et al*., 2021). Intriguingly, they found higher conservation of old-biased genes, a result which would be consistent with the reported absence of reproductive senescence in these ant queens.

This growing body of evidence suggests a role for Medawar’s mutation accumulation process, and specifically, a role for higher drift on late-expressed genes in metazoan aging. Still, caution is warranted. First, the diversity of taxa studied is limited to five species of mammals and three species of insects. Second, studies conducted hitherto have analyzed either bulk tissue (mammals) or whole-body samples (insects). It therefore remains possible that the observed expression trends, and expression-conservation correlations, may in fact be driven by aging-related changes in tissue composition and/or cell type composition within tissues. Indeed, aging causes apparent shifts in cell type composition, including tissue-specific cells and immune cells, as shown in mouse tissues (The Tabula Muris Consortium, 2020), in the human brain (Soreq *et al*., 2017) and in the fruit fly brain (Davie *et al*., 2018). Shifts in a tissue’s cell type composition with age will likewise shift expression patterns measured at the bulk tissue level, even in the absence of cell-autonomous expression change. Furthermore, tissues are known to vary in the average conservation levels of the genes they express (Khaitovich *et al*., 2005), and we may reasonably expect that cell type transcriptomes likewise vary in their average conservation levels. Accordingly, the observed ADICT pattern can have two, non-mutually exclusive explanations: (a) late-expressed genes at the cell-autonomous level being subject to stronger drift, (b) weakly conserved cell types increasing their representation within tissues at late age.

Here we address these issues, first by studying the prevalence of ADICT across a diverse range of organisms using published bulk tissue transcriptome profiles (as opposed to whole organism transcriptomes). Second, we investigate conservation level differences among mouse cell types, and we test whether ADICT can be observed at the cell type-specific level.

## Methods

### Conservation metric

When available, *dN* (nonsynonymous substitution rate) and *dS* (synonymous substitution rate) values were downloaded from the Ensembl BioMart using the most recent Ensembl releases that included these metrics (Ensembl v.99 for *dN* and *dS* values between *G*.*gallus*-*M*.*gallopavo* and *M*.*musculus*-*R*.*norvegicus*; Ensembl Metazoa v.45 for *dN* and *dS* values between *D*.*melanogaster*-*D*.*simulans*) (Yates *et al*., 2019). Only one-to-one orthologs, estimated by Ensembl, were included in the study. The conservation metric was calculated as *-ln(dN/dS)*, with higher values corresponding to higher sequence conservation. Genes with *dN/dS* ≥ 0.8 were excluded to limit the influence of positively selected genes on downstream analysis. Genes with *dN* = 0 and *dS* = 0 were also excluded to avoid zero and infinite *dN/dS* values, respectively. *dN* and *dS* values for killifish gene orthologs were not available in BioMart and we used *dN/dS* values calculated for the “FKK-branch” by Sahm and colleagues (Sahm *et al*., 2017). Statistics related to the conservation metric data (mean and median *dN/dS* values, gene numbers, divergence between species pairs) are provided in Table S1. We also ignored genes with paralogs in this analysis in order to limit uncertainty and ambiguity when calculating dN/dS ratios.

### Relative conservation score

To simplify interpreting and comparing the conservation score values when visualizing, we scaled these using the conservation scores of constantly expressed genes (defined below). For this, in each dataset, we calculated a “relative conservation score” for each gene *i* as:

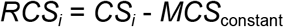

where *CS*_*i*_ is the protein sequence conservation score for that gene (as described above), and *MCS*_constant_ is the arithmetic mean conservation score for all genes classified as constantly expressed in that dataset.

### Bulk RNA-seq data analyses

#### Pre-processing

Raw FASTQ files for the mice brain (SRP108790, GSE99791) (Boisvert *et al*., 2018), chicken brain (SRP144776, GSE114129) (Xu *et al*., 2018), and naked-mole-rat brain (SRP007398, GSE30337) (Kim *et al*., 2011) transcriptomes were downloaded from the European Nucleotide Archive (ENA) repository. The dataset contents are listed in Table 1. FASTQ files were assessed for quality using *FastQC* tool (v0.11.9) and trimmed for low quality and adapter contamination using *Trimmomatic* (v0.39) (Bolger *et al*., 2014) with the options *SLIDINGWINDOW:4:15, MINLEN:25* & *<adapter fasta>:2:30:10:8:TRUE*. Trimmed reads were aligned to the reference genome (chicken GRCg6a, mouse GRCm38.p6, NMR HetGla_female_1.0) and counted using *STAR* (v2.7.6a) (Dobin *et al*., 2013) with the parameter *--quantMode GeneCounts*. If present, drug-treated and non-adult samples were discarded from the datasets, along with genes not expressed in any of the remaining samples in each dataset. Resulting count data were normalized using the median ratios method implemented in the DESeq2 (v1.26.0; Love *et al*., 2014) package. Briefy, we used the *DESeqDataSetFromMatrix* function from the DESeq2 package, with the arguments “*design = ∼age”* to construct a DESeqDataSet object, and used the *estimateSizeFactors* function to estimate size factors for normalizing raw counts. For the killifish liver and skin transcriptomes (Reichwald *et al*., 2015) and the fruit fly brain transcriptomes (Pacifico *et al*., 2018), gene count data were used instead of the raw FASTQ files. Count data for these datasets were normalized as before. We studied the pre-processed datasets by performing principal components analyses and plotting the first four principal components (Supplementary Figure 1 & 2).

**Table 1.**
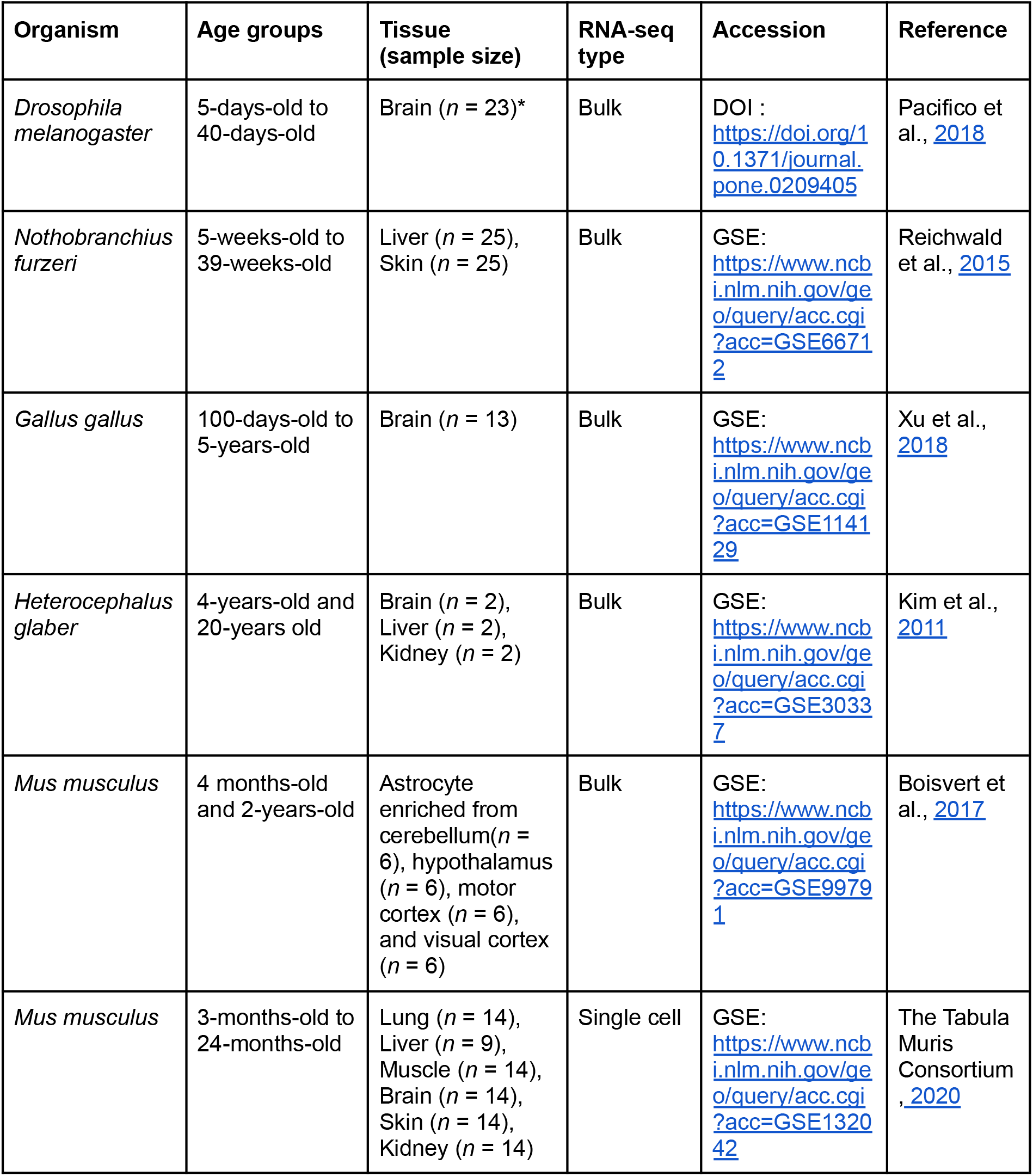
Summary of datasets used in this study. Age groups and tissue sample sizes refer to only those individuals analyzed within this study, i.e. healthy adults (no drug treatment, etc.). (*): each sample represents a pool of 18 individuals.

| Organism | Age groups | Tissue (sample size) | RNA-seq type | Accession | Reference |
| --- | --- | --- | --- | --- | --- |
| *Drosophila melanogaster* | 5-days-old to 40-days-old | Brain ($n = 23$)* | Bulk | DOI : https://doi.org/10.1371/journal.pone.0209405 | Pacifico et al., 2018 |
| *Nothobranchius furzeri* | 5-weeks-old to 39-weeks-old | Liver ($n = 25$), Skin ($n = 25$) | Bulk | GSE: https://www.ncbi.nlm.nih.gov/geo/query/acc.cgi?acc=GSE66712 | Reichwald et al., 2015 |
| *Gallus gallus* | 100-days-old to 5-years-old | Brain ($n = 13$) | Bulk | GSE: https://www.ncbi.nlm.nih.gov/geo/query/acc.cgi?acc=GSE114129 | Xu et al., 2018 |
| *Heterocephalus glaber* | 4-years-old and 20-years old | Brain ($n = 2$), Liver ($n = 2$), Kidney ($n = 2$) | Bulk | GSE: https://www.ncbi.nlm.nih.gov/geo/query/acc.cgi?acc=GSE30337 | Kim et al., 2011 |
| *Mus musculus* | 4 months-old and 2-years-old | Astrocyte enriched from cerebellum($n = 6$), hypothalamus ($n = 6$), motor cortex ($n = 6$), and visual cortex ($n = 6$) | Bulk | GSE: https://www.ncbi.nlm.nih.gov/geo/query/acc.cgi?acc=GSE99791 | Boisvert et al., 2017 |
| *Mus musculus* | 3-months-old to 24-months-old | Lung ($n = 14$), Liver ($n = 9$), Muscle ($n = 14$), Brain ($n = 14$), Skin ($n = 14$), Kidney ($n = 14$) | Single cell | GSE: https://www.ncbi.nlm.nih.gov/geo/query/acc.cgi?acc=GSE132042 | The Tabula Muris Consortium , 2020 |

#### Differential gene expression analysis

Genes that show aging-related expression changes were identified using Spearman correlation: a gene was classified as age-related if the expression of the gene showed a statistically significant correlation with age after multiple testing correction (Benjamini-Hochberg corrected *p* < 0.1), and an absolute expression vs. age correlation coefficient (*rho*) higher than 0.5. Note that we only used data from adult individuals and excluded pre-adults (Table 1). Among age-related genes, genes with positive expression-age correlation were classified as “old-biased”, and the genes with negative expression-age correlation were classified as “young-biased”. Genes outside either class were classified as “constantly expressed”. Genes assigned to each gene class are shown in Table S2. For the naked-mole-rat, due to lack of biological replicates in this dataset, genes with positive Spearman correlations between expression vs. age were classified as old-biased genes and the ones with negative correlation were classified as young-biased genes, without reference to statistical significance.

#### Transcriptome conservation

Here we followed the approach outlined by Turan and colleagues (Turan *et al*., 2019). For each individual sample in a dataset, we calculated a single expression-conservation correlation measure, reflecting the correlation between normalized gene expression values and the conservation metric across all genes, using the Spearman correlation test via the *cor* function in *R* (v. 3.6.3) *stats* package, with the arguments *“method = spearman”*. We then calculated the correlation between individual age and expression-conservation correlations across all individuals in a dataset, again using Spearman correlation. This latter correlation was treated as a measure of age-related change in overall transcriptome conservation.

#### Conservation difference between old-biased and young-biased genes

To test whether the distribution of conservation scores differs between old-biased and young-biased genes we used the Welch’s t-test (two sided) on conservation scores of old and young-biased genes separately for each tissue in each dataset, as implemented by the *t*.*test* function in the *stats* package of *R* with default parameters. As a non-parametric alternative, we also used the Mann-Whitney U test as implemented by the *wilcox*.*test* function in the *R stats* package, using default parameters.

### Single cell RNA-seq data analyses

#### Pre-processing

Single cell expression data for six different tissues (lung, liver, muscle, brain, skin, and kidney) were downloaded from the Tabula Muris Senis dataset (The Tabula Muris Consortium, 2020) and processed following Izgi and co-authors (Izgi *et al*., 2022). For each cell type, the gene expression levels per individual were calculated as the mean expression value across cells of that cell type from a given individual, calculated separately for each gene, and separately in each tissue. We removed cell types absent in any of the three age groups (3-month-old, 18-month-old, 24-month-old). To minimize the individual effect in downstream analyses, we limited our analysis to include only individuals that had expression data for >70% of all cell types present within a given tissue for a given age group (irrespective of the number of cells measured for that cell type). The final number of cell types and individuals are presented in Table S3.

#### Cell type-specific differences in transcriptome conservation

To evaluate the cell type-specific differences in transcriptome conservation, in each tissue we calculated the Spearman correlation coefficient between expression levels and conservation scores as above, only using the three-months old individuals for every cell type. We excluded cell types represented by less than three individuals. We then tested the cell type difference in expression-conservation correlation using ANOVA as implemented in the *aov* function in the *R stats* package with the model “*rho ∼ tissue + cell type*”. We additionally tested the effect of immune status of cell types on transcriptome conservation levels using a mixed model ANOVA via the *lme* function of the *R nlme* package (v.3.1-144) with transcriptome conservation set as the response variable, immune status as the explanatory variable and tissue as a random effect.

#### Differential gene expression analysis

For each cell type in each tissue, genes that showed aging-related differential expression were identified using Spearman correlation between the expression level of the gene vs. individual age (as implemented on bulk RNA-seq datasets). Due to higher noise in the scRNA-seq dataset compared to bulk RNA-seq datasets, a cut-off of rho (⍴) > 0.5 was used to define age-related genes, instead of using both *rho* and *p*-value cut-offs. Age-related genes with *rho* < -0.5 were classified as young-biased genes and those with *rho* > 0.5 were classified as old-biased genes.

#### Transcriptome conservation

Expression-conservation correlations and related analyses were conducted identically to those conducted on bulk RNA-seq data.

#### Conservation difference between old and young-biased genes

To test whether the distribution of relative conservation scores differs between old-biased genes and young-biased genes, we calculated the mean relative conservation score (MRCS) of these gene sets identified per cell type in each tissue (i.e. one estimate per gene set per cell type). Then, across all cell types in each tissue, we applied the Wilcoxon signed rank test on the distributions of MRCS of old-biased and young-biased genes using the *wilcox*.*test* function with the parameter *“paired = TRUE ‘‘* in the *R stats* package.

#### Immune cells

To test whether immune cells behave differently in terms of age-related changes in sequence conservation than non-immune cells, we calculated the difference between mean relative conservation scores of old-biased and young-biased genes for each cell type in the dataset by subtracting the mean relative conservation scores of old-biased genes from the mean relative conservation scores of young-biased genes. The list of cells used and their immune status is shown in Table S4. Next, we tested whether the distributions of MRCS differences and ADICT signals (Spearman correlation between expression-conservation correlation vs. age) differed between immune and non-immune cells. For cell types that are present in multiple tissues, we used the mean of MRCS differences calculated in each tissue as the MRCS difference score for the cell type. We similarly used the mean rho scores across different tissues for repeating cell types when comparing ADICT signals. For the statistical comparison, we used the Mann-Whitney U test as implemented by the *wilcox*.*test* function with default parameters in the *R stats* package. Additionally we calculated effect sizes for the differences using Cohen’s d, for both the MRCS differences and differences in ADICT signals (Spearman correlation distributions).

## Results

### Age-related decrease in transcriptome conservation is observed across diverse metazoan species

We first asked whether the previously described pattern of age-related decrease in evolutionary transcriptome conservation (ADICT) (Turan *et al*., 2019) can also be observed in tissue-specific transcriptomes of non-mammalian organisms. To this end, we collected published aging transcriptome datasets from diverse taxa, including a *G. gallus* (chicken) brain aging dataset, a *N. furzeri* (turquoise killifish) liver and skin aging dataset, and a *D. melanogaster* (fruit fly) brain aging dataset (Table 1).

Using the same framework used by Turan and colleagues (Turan *et al*., 2019), we calculated the correlation between protein sequence conservation metrics (based on *dN*/*dS*; see Methods) and expression levels across genes for each individual. We used this correlation value as a metric of transcriptome conservation which integrates expression levels of genes in the transcriptome. We then looked at how transcriptome conservation changes with age by studying the relationship between the expression-conservation correlation scores and individual age (Figure 1A, showing the *G. gallus* brain dataset as example). The results for the whole transcriptome revealed moderate ADICT patterns in all four datasets, i.e. negative correlations between expression-conservation correlation and age (Figure 1B, summarized in Table S5). The ADICT signal became more conspicuous when we limited the analysis to differentially expressed genes (Figure 1A-B).

**Figure 1.**
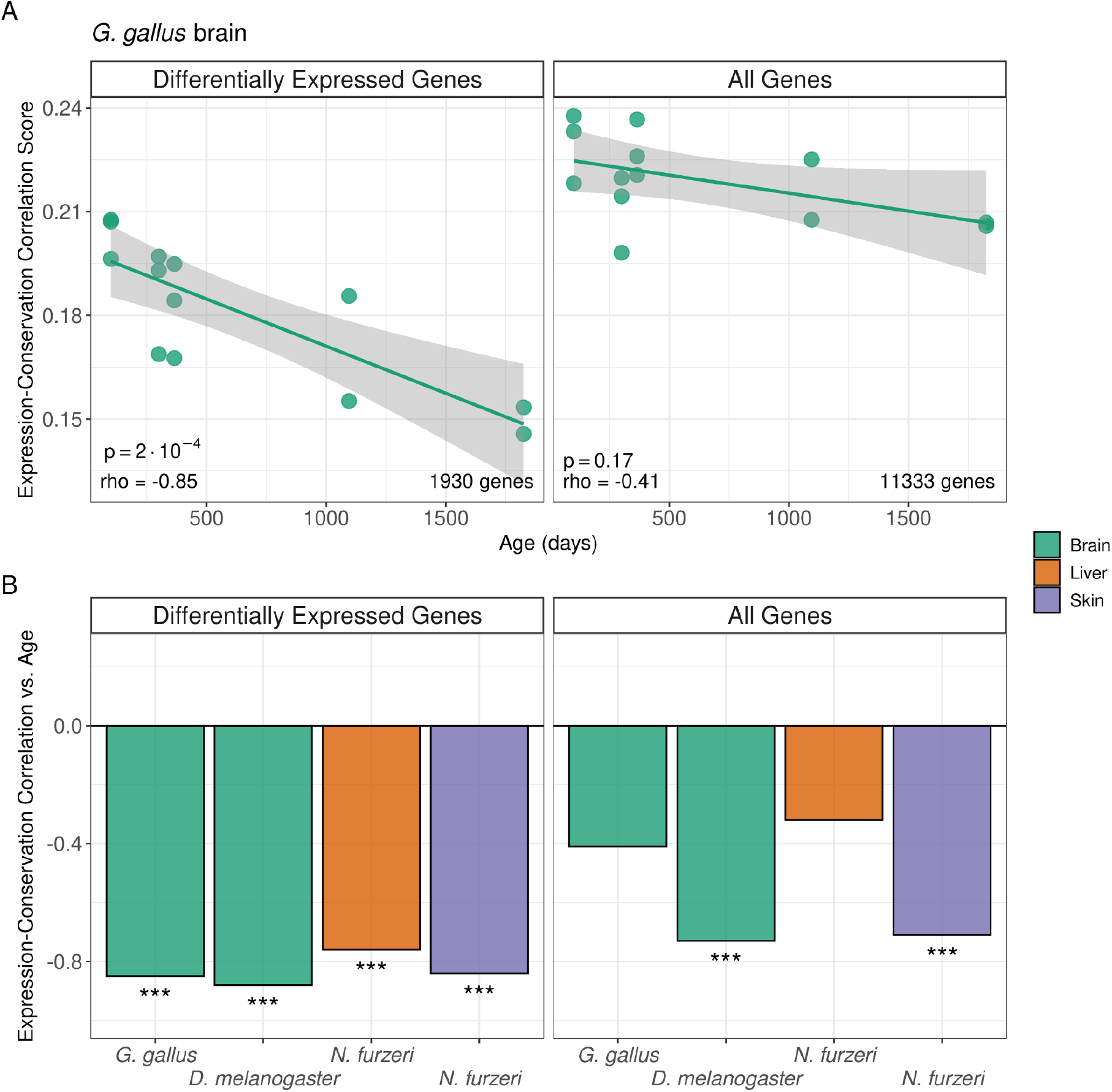
Age-related changes in expression-conservation correlation in diverse metazoa. (A) Age-related changes in expression-conservation correlation in the *G. gallus* brain. The y-axis shows the Spearman correlation coefficient between expression level and the protein sequence conservation metric across genes for each individual in this dataset. Each dot corresponds to an individual (*n* = 13). The x-axis shows the individual age. The left panel shows the analysis result using genes differentially expressed with age (*n* = 1930), and the right panel shows the result using the whole transcriptome (*n* = 11,333). Spearman correlation *rho* and p-values in the inset are calculated between the expression-conservation correlation and individual age. (B) Barplot of Spearman correlation values between the expression-conservation correlation vs. age, (***): *p* < 0.001. Tissues are illustrated in different colors (key shown on the right hand side).

We then asked whether the overall negative correlations are driven by highly conserved genes decreasing in expression with age or by weakly conserved genes increasing in expression with age, or both. To this end, we examined conservation levels of young-biased genes (that show significant negative expression vs. age correlation with rho < -0.5) and old-biased genes (that show significant positive expression vs. age correlation with rho > 0.5) for each data set. Old-biased gene sets showed consistently lower average conservation than young-biased genes in all four datasets (Welch’s t-test *p*<0.003 across all four tests, Figure 2A). Old-biased and young-biased gene sets also respectively showed trends of lower and higher conservation compared to constantly expressed genes (Figure 2B). Turan and colleagues had also observed similar trends, with some degree of variability across mammalian tissues (Turan *et al*., 2019).

**Figure 2.**
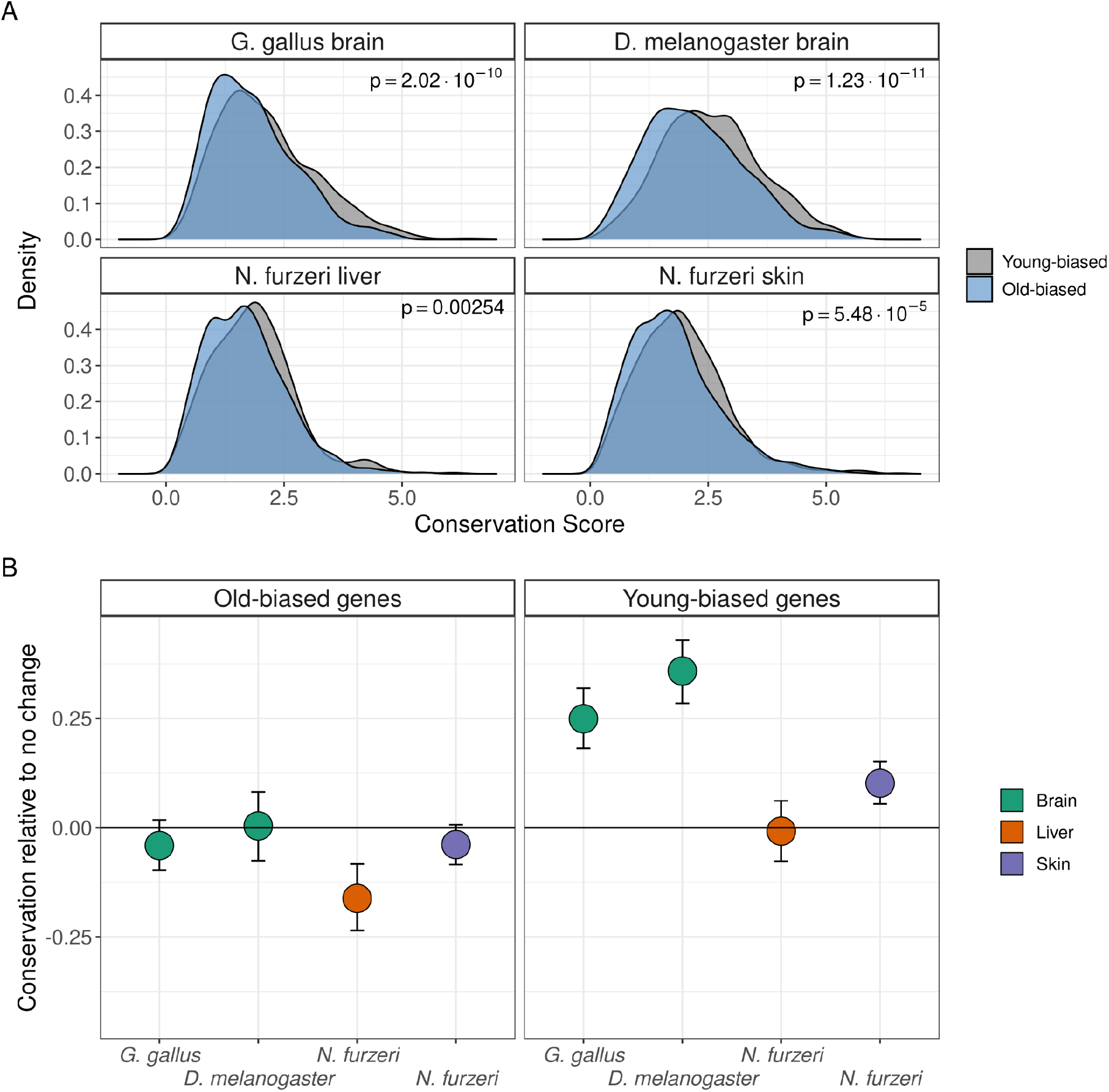
(A) Density plots of conservation scores of old-(blue) and young-biased (gray) genes, relative to constantly expressed genes. P-values indicate the results of Welch’s *t*-test (two-sided) between the distributions of old-biased and young-biased genes. (B) Mean conservation scores of old-biased and young-biased genes, relative to constantly expressed genes. Error bars indicate 95% confidence intervals calculated using 1,000 bootstraps (*D. melanogaster* brain *n*_*old-b*._= 792 genes, *n*_*young-b*._= 831 genes; *G. gallus* brain *n*_*old-b*._= 1066 genes, *n*_*young-b*._= 864 genes; *N. furzeri* skin *n*_*old-b*._= 1468 genes, *n*_*young-b*._= 1405 genes; *N. furzeri* liver *n*_*old-b*._= 537 genes, *n*_*young-b*._= 618 genes).

In addition, we analyzed published transcriptome data from naked mole rats (NMR) *H. glaber*, an exceptionally long-lived eusocial rodent, which can survive more than 30 years in captivity (Kim *et al*., 2011). Both breeding and non-breeding NMR were recently reported not to show increasing age-specific hazard of mortality (Ruby *et al*., 2018), consistent with the case of long-lived ant queens described earlier (Harrison *et al*., 2021). The dataset consists of tissue samples from brain, kidney and liver of a single 4-year-old and another 20-year-old female NMR. Consequently, we could only classify genes into old-biased and young-biased groups based on expression trends. In all three tissues, old-biased genes showed weaker conservation than young-biased genes, although we note that the lack of biological replicates does not allow generalizing the result (Supplementary Figure 3).

### Cell type transcriptomes vary dramatically in average conservation levels

We reasoned that ADICT patterns (lower conservation of late-expressed genes) we observed on bulk-tissue transcriptomes could be driven by (a) cell type composition changes, such that cell types with weakly conserved transcriptomes become more abundant in tissues or (b) cell type specific changes, such that late-expressed genes in each specific cell type tends to be weakly conserved, or both. Notably, scenario (a) assumes the presence of significant differences among specific cell types with respect to average conservation levels of their transcriptomes. To investigate such possible conservation differences among cell-types, we used published single cell transcriptome data from 44 cell types across lung, skeletal muscle, brain, skin, and kidney (The Tabula Muris Consortium, 2020). We estimated a transcriptome conservation level for each cell type, calculated as the correlation between conservation scores and the mean gene expression levels of all cells assigned to that cell type in a given individual, using young adult mice (Methods).

The results shown in Figure 3 demonstrate salient variation among specific cell types in their average transcriptome conservation levels. There was a significant cell type effect along with a tissue effect in a two-way ANOVA (F_tissue_= 134.22, d.f_tissue_= 4, F_cellType_= 33.03, d.f_cellType_= 43, *p*<1×10^−15^ for both factors; Table S6). In general, neural cells, oligodendrocytes and astrocytes showed the highest transcriptome conservation, parallel to earlier observations indicating high protein sequence conservation of brain-specific genes (Khaitovich *et al*., 2005). Certain specialized and proliferative cells in other tissue types, such as keratinocytes, skeletal muscle satellite cells and mesenchymal stem cells also showed higher-than-average transcriptome conservation. Immune cells showed the opposite trend, with various types (including B-cells, T-cells, neutrophils, myeloid dendritic cells) on the lowest end of the transcriptome conservation distribution in our sample. A mixed model ANOVA with immune status as the explanatory variable and tissue as random effect also revealed a negative significant effect of immune status on transcriptome conservation (F_immuneStatus_= 95.79, d.f = 1, *p*<0.0001). However, immune cells were not alone in low transcriptome conservation patterns; some tissue-specific cell types, such as the epithelial cell of the proximal tubule (kidney) and the club cell of the bronchiole (lung) also displayed low transcriptome conservation. We also repeated the same analysis using cell type transcriptomes of 18-month-old and 24-month-old mice, which revealed similar results (Figure S4).

**Figure 3.**
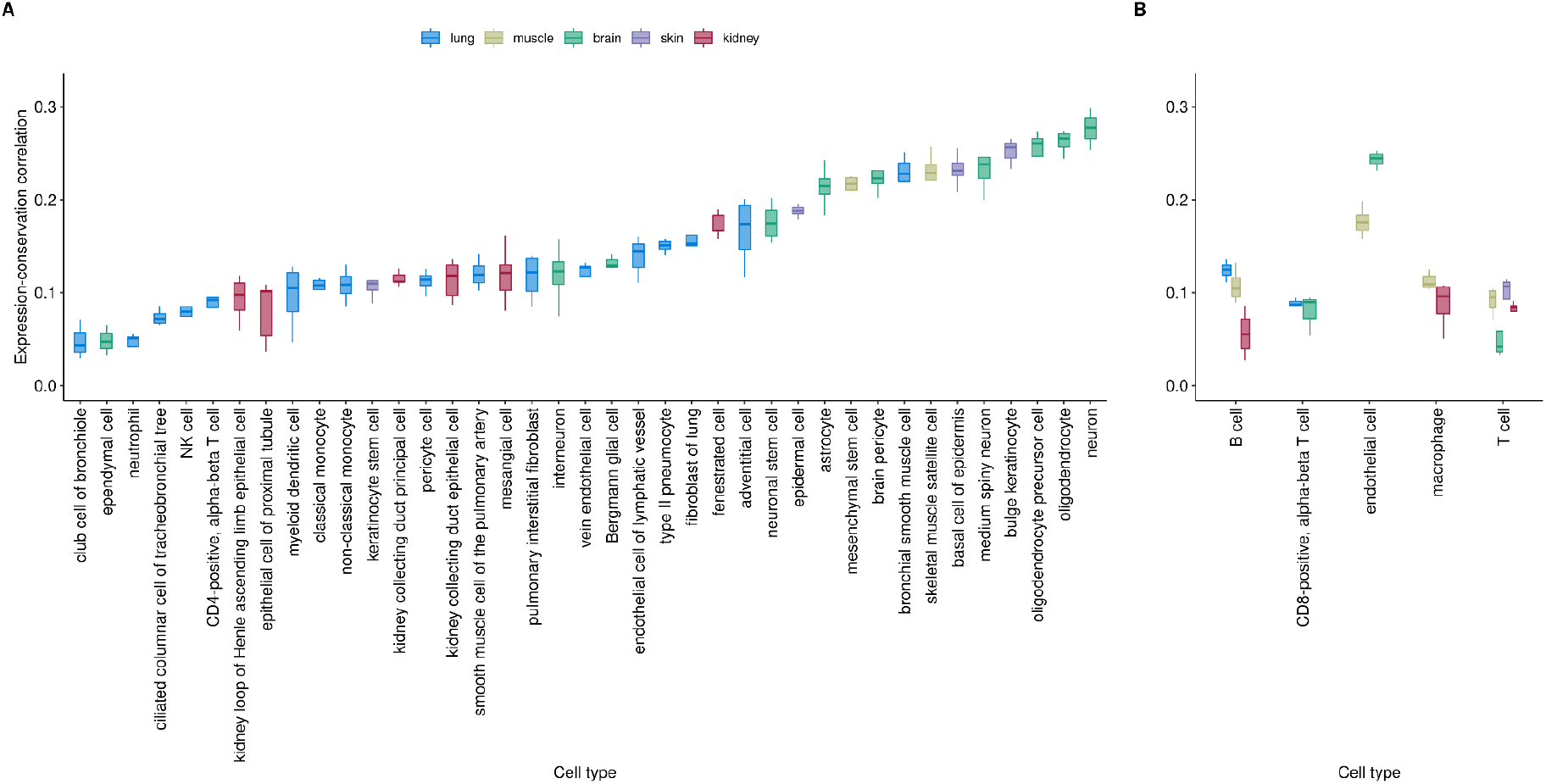
Variability of expression-conservation correlation levels across cell-type transcriptomes of young-adult (3-month-old) individuals. Each data point within bar plots represents Spearman’s correlation values for an individual, calculated using gene expression values averaged across cells belonging to that individual. Panel A and B show cell-types only sampled in one tissue and cell-types sampled from multiple tissues, respectively. Color-coding indicates the tissues which the cell-types were sampled from (see key at the top of the figure). Cell types with less than three correlation values (i.e. individuals) are excluded.

### Cell type-specific changes in transcriptome conservation with age

The strong variation in transcriptome conservation observed among mammalian cell types raises the possibility that cell type composition changes might be the sole driver behind ADICT. If true, we should observe no ADICT signal *within* cell type-specific transcriptomes. To address this, we leveraged upon two cell type-specific aging transcriptome datasets. The first was an aging transcriptome dataset (Table 1) consisting of young (4-month-old, *n* = 3) and old (24-month-old, *n* = 3) mouse tissue samples from cerebellum, hypothalamus, motor cortex, and visual cortex; enriched for astrocytes using astrocyte-ribotagging, which is able to differentially capture actively translated portions of the transcriptome from tagged astrocytes (Boisvert *et al*., 2018). From four different brain regions, we identified significant age-related expression changes (see Methods) in the cerebellum and hypothalamus, which we investigated further. Analyzing the data using the same framework described earlier, we found a negative correlation between expression-conservation and age using differentially expressed genes in astrocyte transcriptomes of the cerebellum and hypothalamus (Figure 4 and S5; Table S5).

**Figure 4.**
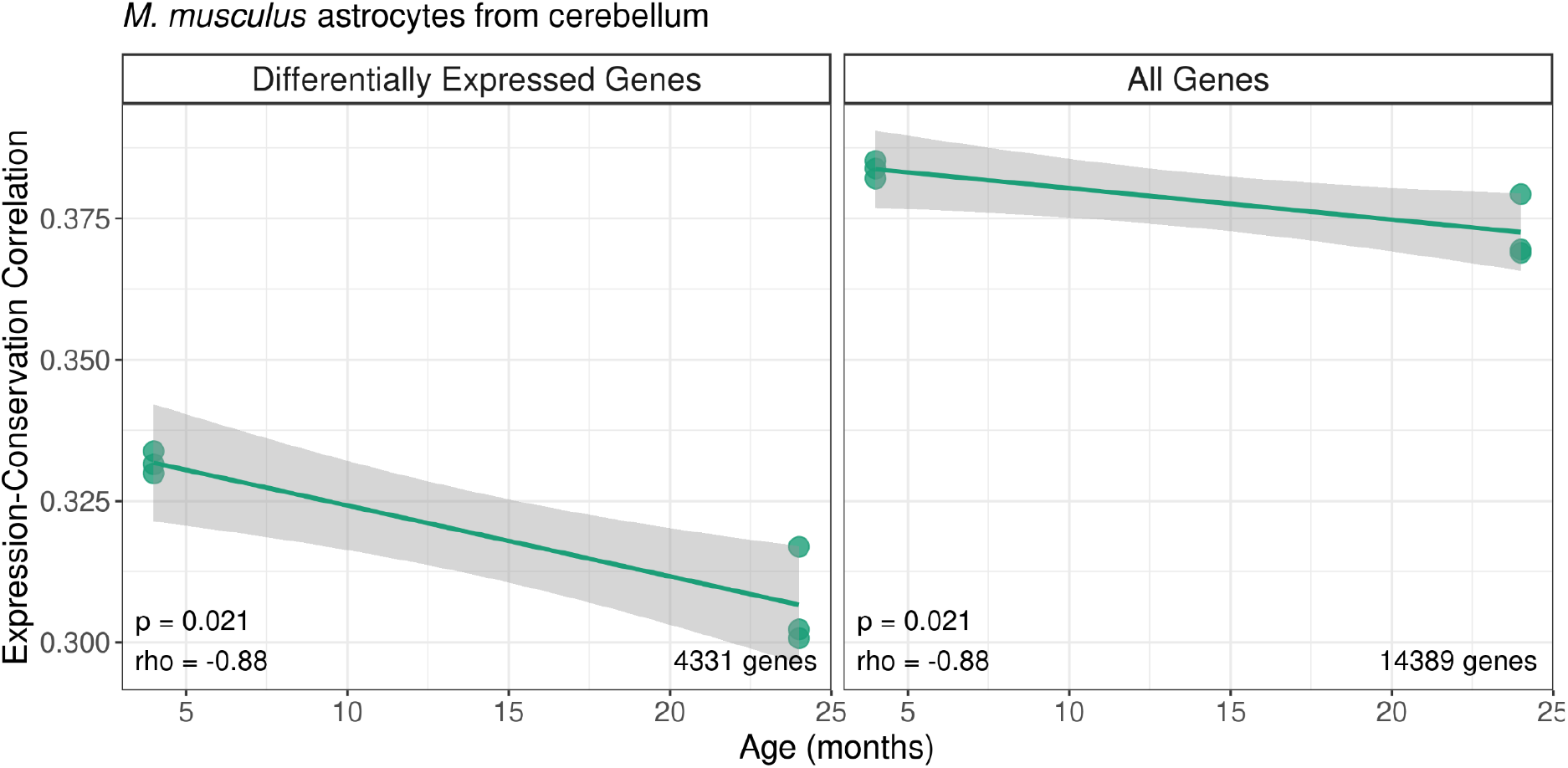
Age-related changes in expression-conservation correlation in *M. musculus* astrocyte transcriptome from the cerebellum. The y-axis shows the Spearman correlation coefficient between expression level and protein sequence conservation metric across genes for each individual in this dataset (*n* = 6). The x-axis shows individual age. The left panel shows the analysis results using genes differentially expressed with age (*n* = 4,331), and the right panel shows the analysis results using the whole transcriptome (*n* = 14,389). *rho* and p-values in the inset indicate the results of the Spearman correlation between the expression-conservation correlation and age. A Mann-Whitney U test on the same data yields a p-value of 0.1.

We next analyzed single-cell transcriptomes from the Tabula Muris Senis dataset, comprising a total of 23,538 cells across 53 cell types from 14 different individuals, covering 3-months to 24-months of age (*n*_*lung*_ = 12, *n*_*liver*_ = 7, *n*_*muscle*_ = 14, *n*_*brain*_ = 9, *n*_*skin*_ = 10, *n*_*kidney*_ = 14) (The Tabula Muris Consortium, 2020) (Table S3). Among the cell types that are not shared between tissues, 32 out of 44 showed the ADICT pattern, i.e. negative correlations between expression-conservation correlation and age, of which six cell types were significant at Spearman correlation test *q*<0.05, and eight were significant at *q*<0.1. Among the nine cell types that were detected in more than one tissue, five (CD4-positive alpha-beta T cell, CD8-positive alpha-beta T cell, mature NK T cell, macrophage, and endothelial cell) showed consistent ADICT signal across the tissues, three (T cell, NK cell, and B cell) showed inconsistent signals, while one (neutrophil) showed a positive correlation (Figure 5). Only in the skin did no cell type reach this significance threshold (Figure 5). Across all tissues, only one cell type, lung neutrophil, showed positive correlation that reached statistical significance at *q*<0.1 (Figure 5, Table S7).

**Figure 5.**
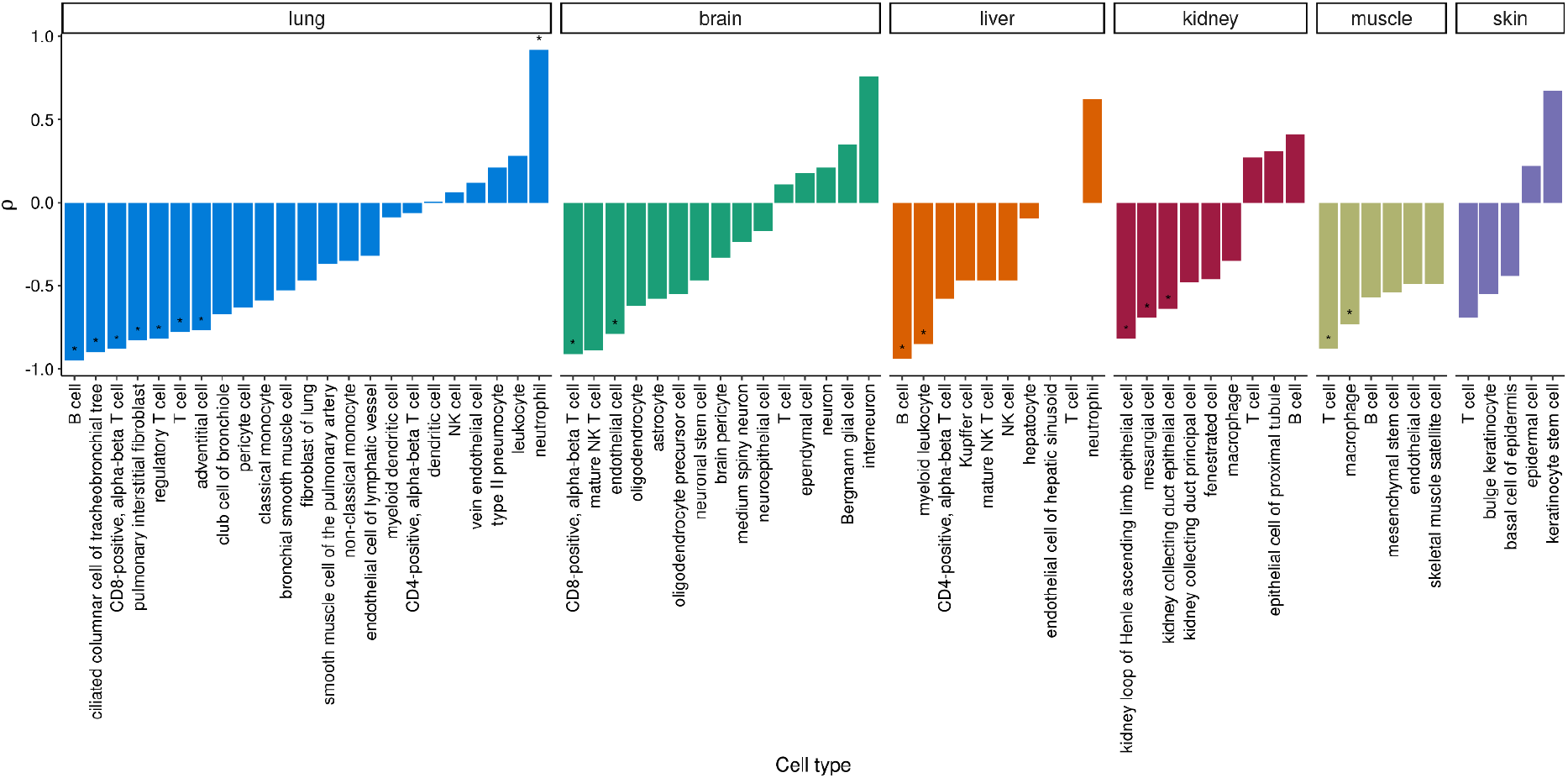
Spearman correlation results for expression-conservation correlation (for differentially expressed genes) vs. age in six different tissues from the Tabula Muris Senis dataset. (*) marks indicate statistical significance at BH corrected p-value <0.1.

We further compared average conservation between old-biased and young-biased gene sets identified in each of the cell types separately for each tissue. Across cell types in all six tissues we observed a trend towards lower conservation among old-biased genes relative to young-biased genes identified in each cell type; this was significant in four tissues (two-sided Wilcoxon signed rank test *p*<0.05) (Figure 6). ADICT can thus be observed at the cell type specific transcriptome level, at least in the mouse and for a substantial number of cell types. Meanwhile, the signal is heterogeneous both among tissues and also among cell types within a tissue.

**Figure 6.**
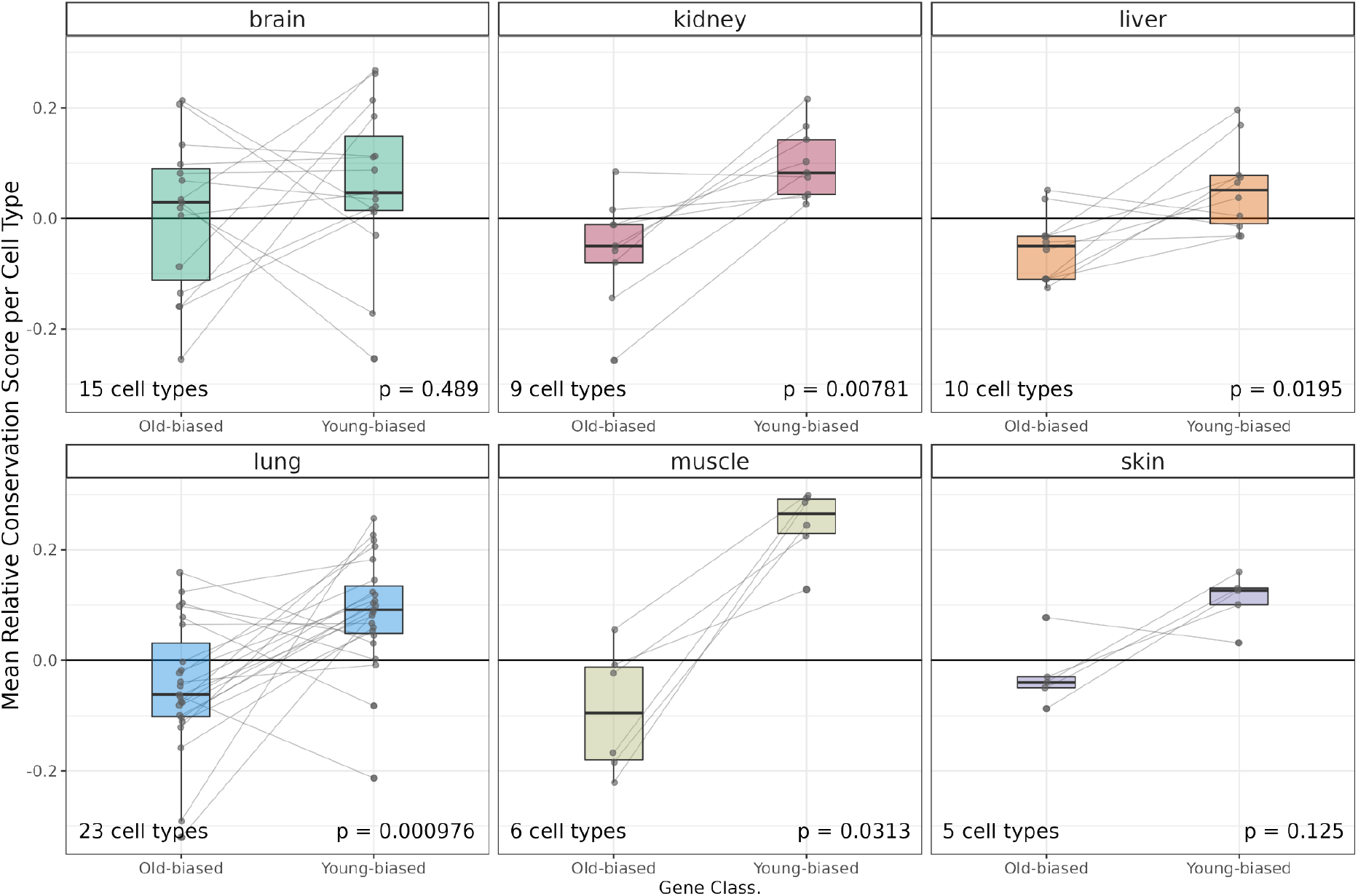
Conservation differences between old- and young-biased genes per cell type. The y-axis shows the mean relative conservation score (relative to constantly expressed genes; see Methods) for old-biased and young-biased genes in each cell type, in each tissue (sample sizes for cell types are n_brain_ = 15, n_kidney_ = 9, n_liver_ = 10, n_lung_ = 23, n_muscle_ = 6, n_skin_ = 5 respectively). The lines connect the mean relative conservation values calculated for old-biased and young-biased gene sets for the same cell type. The p-values indicate Wilcoxon signed rank test results for the difference between the distribution of per cell type mean conservation values of old-biased and young-biased genes in that tissue (no correction for multiple testing).

### ADICT propensities of immune versus non-immune cells

Dysregulated immune responses and inflammation are among the most common hallmarks of aging (López-Otín *et al*., 2013). Aging-related chronic inflammation is known to result in an increase in immune cell types within inflamed tissues. Immune cell types were also ubiquitous across the mouse tissues we analyzed here. We have earlier shown that immune cells have lower conservation levels than average. Here we further asked whether stronger ADICT propensity among immune cell types might be a contributor of ADICT patterns at the bulk-tissue level. Specifically, we tested whether immune cell types show a difference in ADICT propensity compared to other cell types using 16 immune and 37 non-immune cell types from the Tabula Muris Senis dataset (Table S4). For this, (a) we compared ADICT signals between immune and non-immune cell types using Spearman’s rho values between conservation-correlation vs. age, and (b) calculated the mean relative conservation score (MRCS) differences between young-biased and old-biased genes for immune and non-immune cell types, and compared the two (see Methods). Neither analysis revealed a significant difference between immune and non-immune cell types (Mann-Whitney U test *p*-values = 0.83 and 0.69, Cohen’s d = 0.105 and 0.177 for (a) and (b), respectively) (Figure 7).

**Figure 7.**
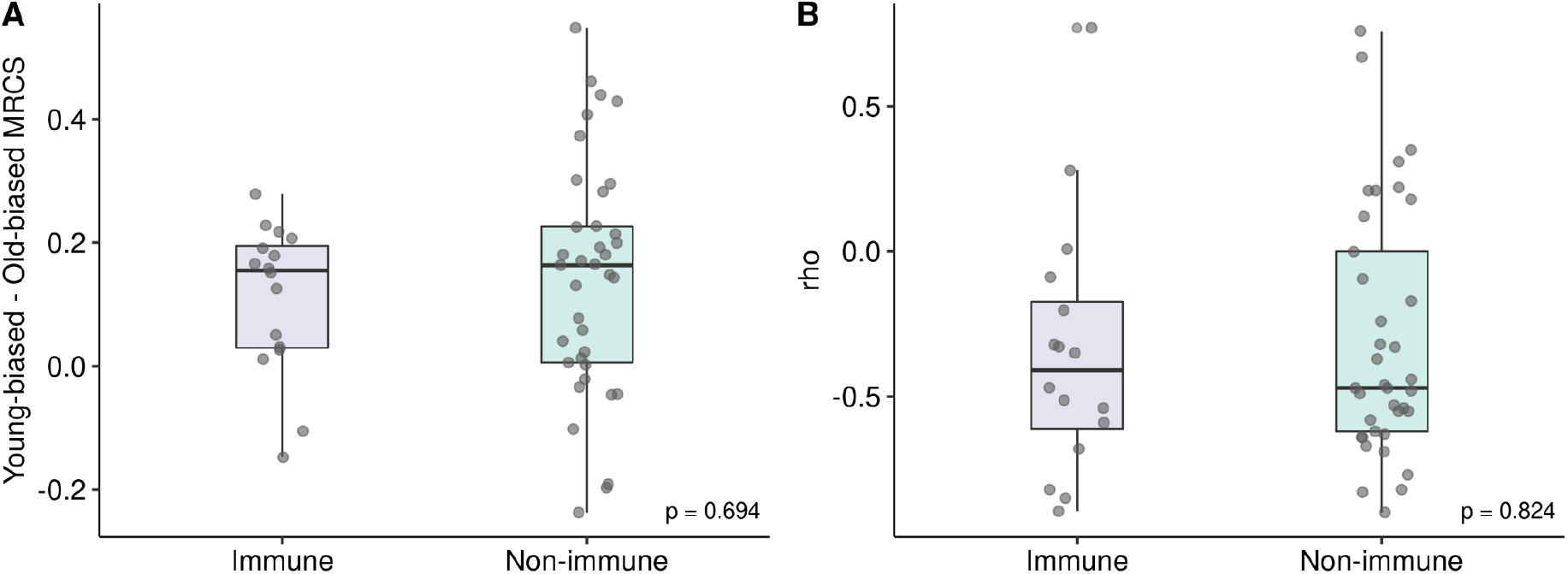
Distributions of (A) mean relative conservation score differences between decreasing and increasing genes and (B) Spearman’s correlation values for expression-conservation correlation vs. age for immune and non-immune cell types. P-values indicate results of a Mann-Whitney U test.

## Discussion

Our results indicate ADICT as a common phenomenon across diverse metazoan taxa with classical senescence patterns, i.e. increased mortality with age. Lower conservation of old-biased genes was previously reported in primate and rodent tissues (Somel *et al*., 2010; Jia *et al*., 2018; Turan *et al*., 2019), as well as whole body transcriptomes of Diptera (Cheng and Kirkpatrick, 2021). We now add fish, birds and fruit flies to the list of taxa in which ADICT has been observed at the tissue level. Interestingly, we also observe low sequence conservation among old-biased genes in three tissues of the extreme long-lived naked mole rat, although the lack of biological replication in this dataset limits our ability to interpret this result. This supports a widespread role of the selection shadow in shaping organismal senescence.

We further show that average transcriptome conservation levels are highly variable between different cell types, at least in the mouse. This result echoes observations on varying conservation levels among mammalian tissues (Khaitovich *et al*., 2005; Kryuchkova-Mostacci & Robinson-Rechavi, 2015). Neurons show highest transcriptome conservation among the 44 mouse cell types analyzed here, which parallels strong conservation of brain-expressed genes observed in these earlier studies, possibly due to sensitivity of neural cells to proteotoxicity (Drummond & Wilke, 2008). Immune cells tend to show weak conservation, which may be partly attributable to the relatively high rates of positive selection pressure on immune-related genes [e.g. Chimpanzee Genome 2005]. Although these results are not surprising, they point to the difficulty in interpreting aging-related transcriptome changes and ADICT signals at the bulk tissue and/or whole body level, given aging-related changes in cell type composition with age [e.g. (The Tabula Muris Consortium, 2020)].

Our third main finding is that ADICT may prevail at the cell type level at least as widely as at the bulk tissue level. This we show using enriched astrocyte transcriptomes, as well as single-cell RNA sequencing data from six tissues in mice. Importantly, here we are assuming that cell type identities are accurately defined and not affected by aging. With this note of caution, and the fact that we have analyzed single cell data from only a single species, our results suggest that previously identified ADICT patterns are not solely driven by cell type composition changes. We thus believe that the evidence we present here, together with earlier work, marks a major presence of ADICT in metazoan aging. However, two main questions remain.

The first involves the causal role of ADICT in organismal senescence. This can be addressed by selection experiments, by comparing species pairs which have recently evolved differences in lifespan, as well as by studying species with constant or decreasing mortality with age (Jones *et al*., 2014; Cohen, 2018). We predict that the evolution of a constant or decreasing mortality curve, or longevity, should lead to increasing purifying selection on late-expressed genes, which should be measurable using standard transcriptome experiments.

This has been exactly what Harrison and colleagues have observed: higher protein sequence conservation among old-biased genes in ant queens (Harrison *et al*., 2021). However, it remains possible that age-related changes in cell type composition (e.g. increasing proportion of reproductive tissue) could underlie the observed signal. Intriguingly, three tissues of the NMR that we analyze here show expression patterns consistent with ADICT, but the data suffers from lack of biological replication. We believe that continuing this line of work with improved sampling could be highly illuminating with respect to the causal role of mutation accumulation in the evolution of aging. We also note that the presence of ADICT in model organisms in aging research, including the mouse, killifish and fruit fly, opens up the possibility of studying ADICT in longevity selection experiments.

The second question relates to the nature of old-biased and low conserved genes, which needs systematic dissection. This is especially interesting in light of the results by Jia and colleagues (Jia *et al*., 2018) and by Cheng and Kirkpatrick (Cheng & Kirkpatrick, 2021), who showed that late-expressed genes tend also to be evolutionarily younger genes. This brings up the possibility of two distinct mechanisms behind ADICT:

1. ADICT by regulatory drift: Part of metazoan proteomes consists of genes that have relatively recently evolved, are expressed at low levels, and which may be less optimized in their structure and functions, including a propensity towards proteotoxicity. Genetic drift on regulatory sequences and regulatory interactions may drive late-expression among such low conserved genes, as their early expression would not be tolerated. Recruitment of such genes in late adult transcriptomes could drive ADICT and shape the aging phenotype. This would be in line with “early-life inertia” model of aging (e.g. de Magalhaes and Church 2005; Carlsson et al. 2021), as well as empirical evidence for faster evolution of promoter sequences in late-expressed genes in mammals (Turan *et al*., 2019).
2. ADICT by drift on coding sequences: Old-biased genes may have critical roles at old age (e.g. protection against accumulating somatic damage). However, drift on their coding sequences may prevent them from functioning at the highest efficiency. This would then allow processes such as somatic damage accumulation, and consequent aging phenotypes. This latter model would predict some degree of conservation of late age expression patterns.

Noting that the two models are non-mutually exclusive, we believe that uncovering their exact contributions to the selection shadow in aging would be an attractive endeavor.

## Author Contributions

NY, ZGT, Hİ and MS conceived the study, MS and ZGT supervised the study, NY performed analyses with support from ZGT, Hİ, FR and UBA; NY, ZGT, Hİ and MS wrote the manuscript, all authors read and approved the manuscript.

## Acknowledgments

We are grateful to Kıvılcım Başak Vural for their support and help, Alexei Maklakov and Handan Melike Dönertaş for helpful comments on the manuscript, and the METU Compevo Group for discussions. ZGT was supported by a YÖK PhD fellowship.

## Data Availability Statement

Only publicly available datasets were used in this study. All relevant accession numbers are provided in Table 1 and throughout the paper.

## Code Availability

The computer code used in the study are available on https://github.com/NisanYildiz/selection-shadow/

## Supplementary Figures

**Figure S1.**
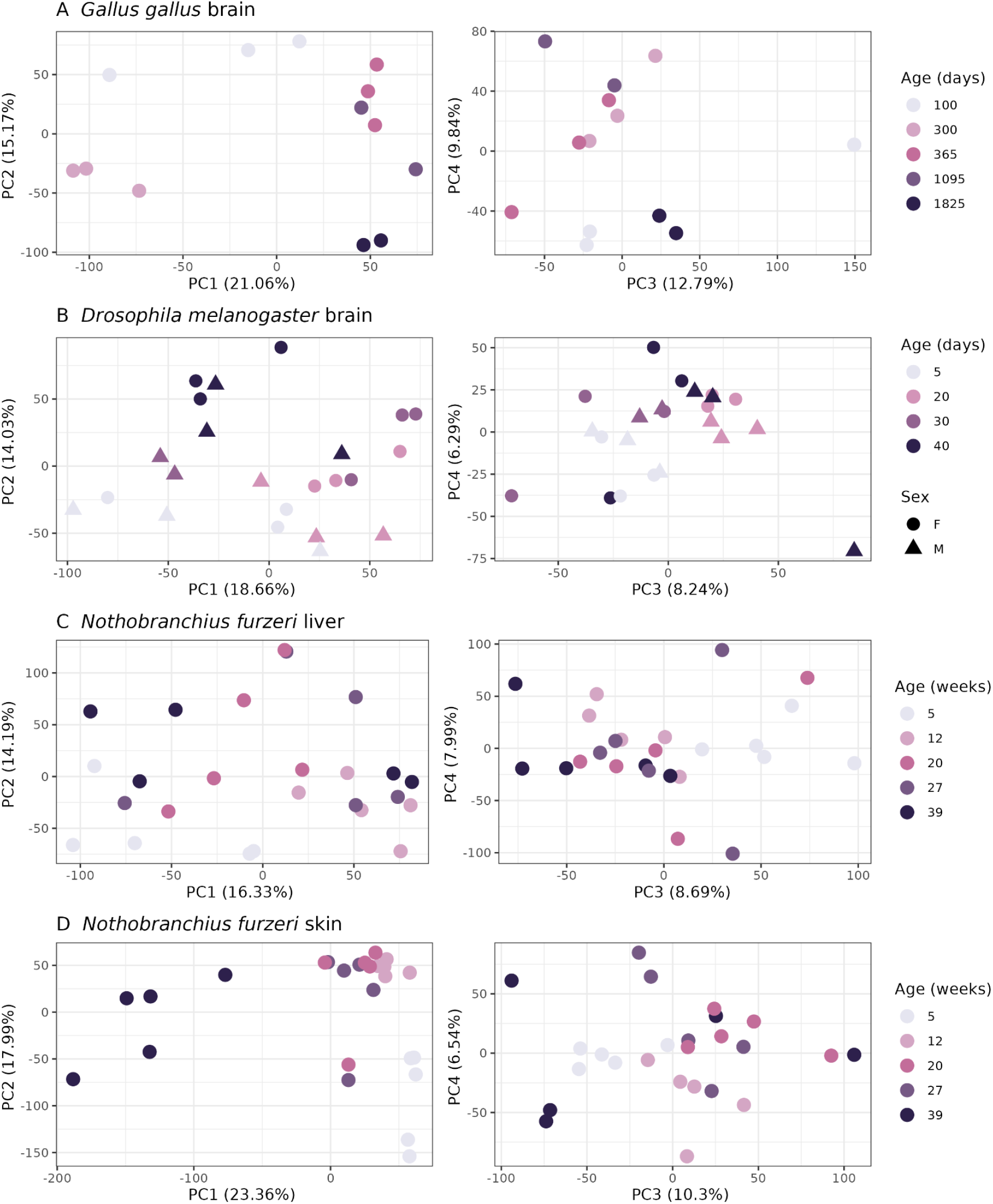
Principal component analysis (PCA) using expression levels of each dataset (*n* = [22405, 12344, 23064, 23178] genes for *G. gallus, D. melanogaster, N. furzeri* liver and *N. furzeri* skin, respectively). Only the first four PCs are plotted. Numbers in the parentheses show the percentage of variation explained by each PC. Age and sex (exist only for *D. melanogaster*) labels are indicated on the right of the plots.

**Figure S2.**
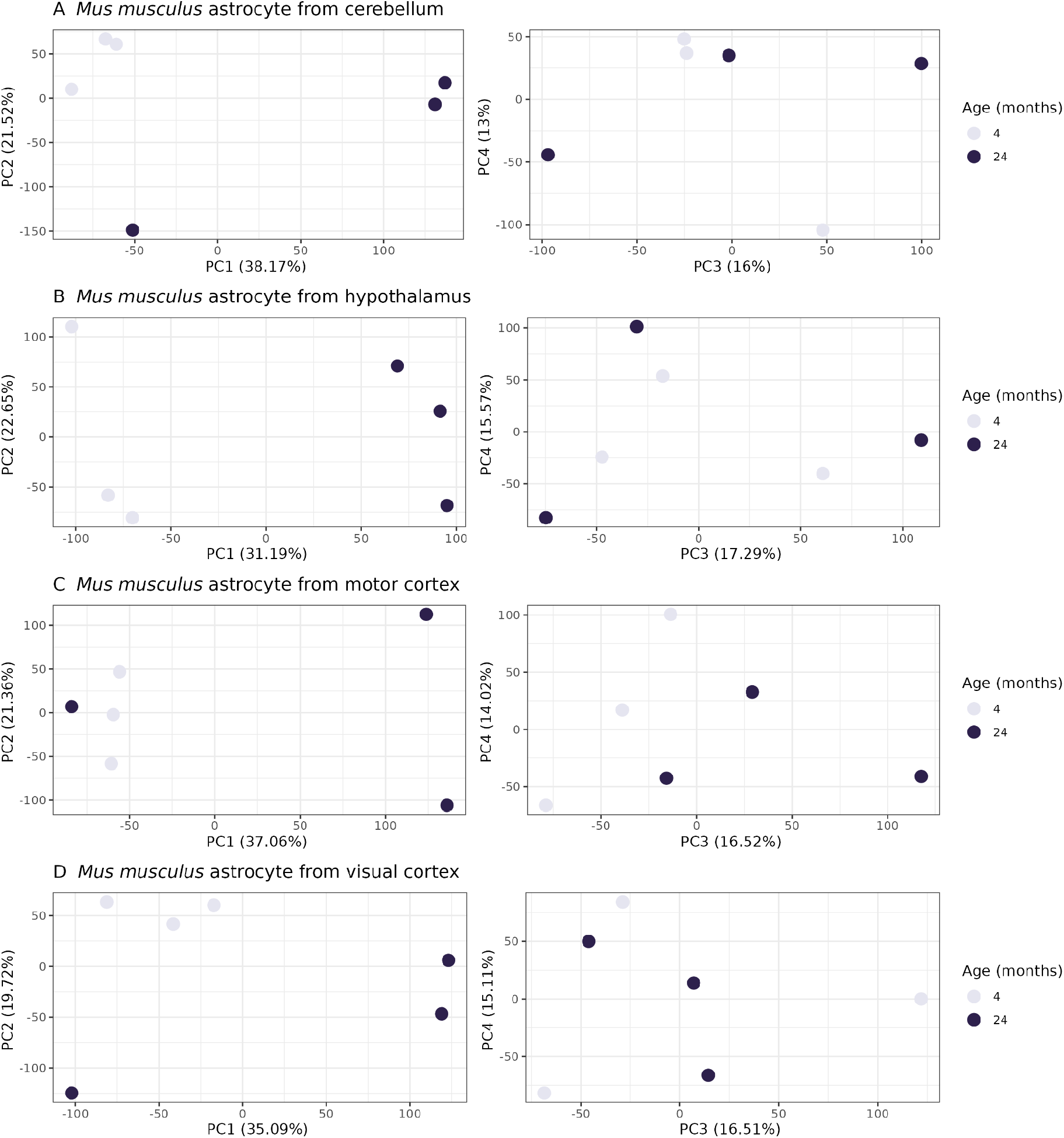
Principal component analysis (PCA) using expression levels of astrocyte-enriched *M. musculus* brain region samples from GSE99791 dataset (*n* = [28572, 28516, 27649, 27580] genes for cerebellum, hypothalamus, motor cortex and visual cortex, respectively). Only the first four PCs are plotted. Numbers in the parentheses show the percentage of variation explained by each PC. Age labels are indicated on the right of the plots.

**Figure S3.**
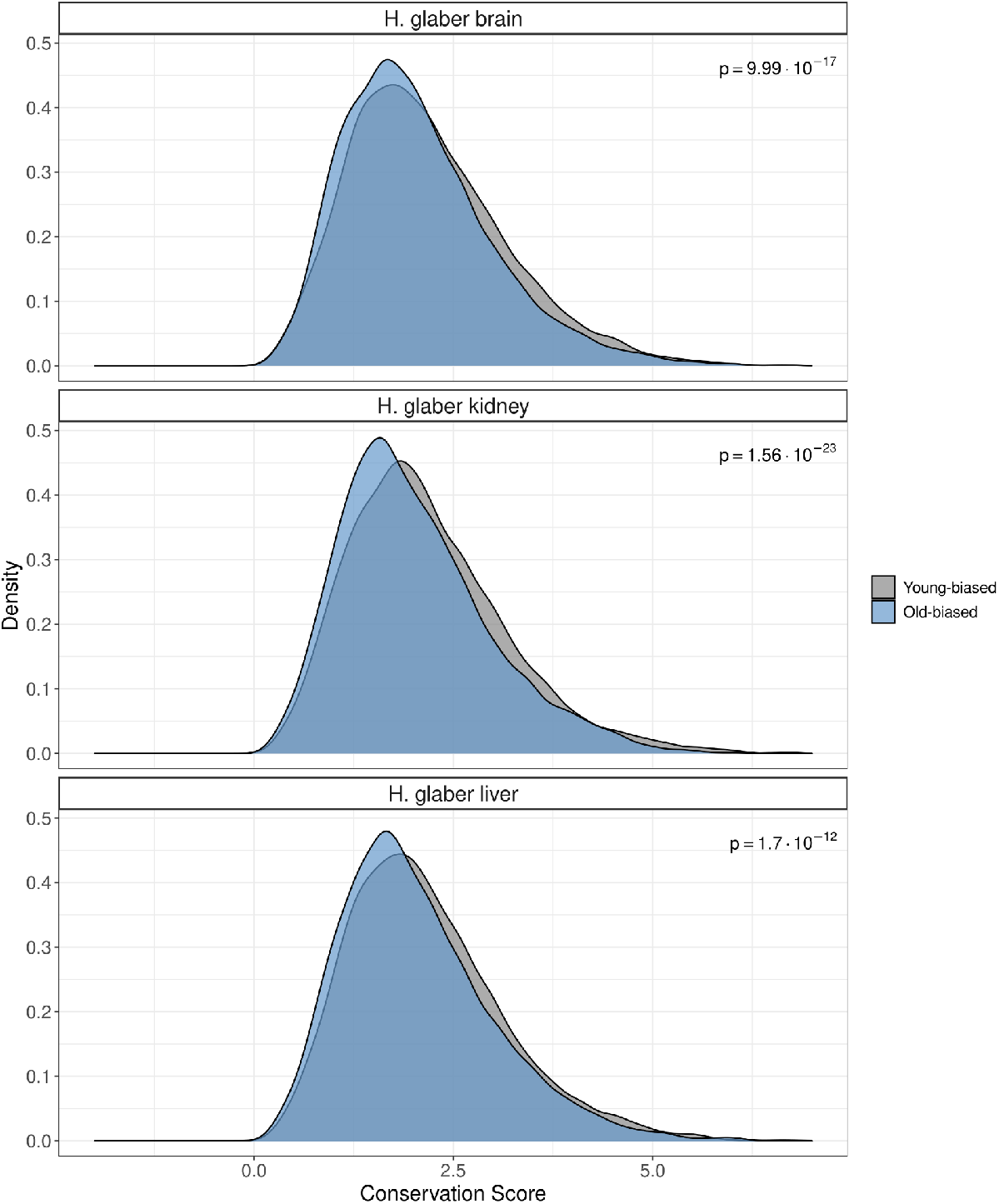
Distribution of the conservation scores (*dN*/*dS*) for young-biased (gray) and old-biased (blue) genes in brain, kidney and liver of *H. glaber*. Old-biased genes have consistently lower conservation scores compared to young-biased genes across tissues. P-values indicate Welch’s *t*-test results between young- and old-biased genes.

**Figure S4.**
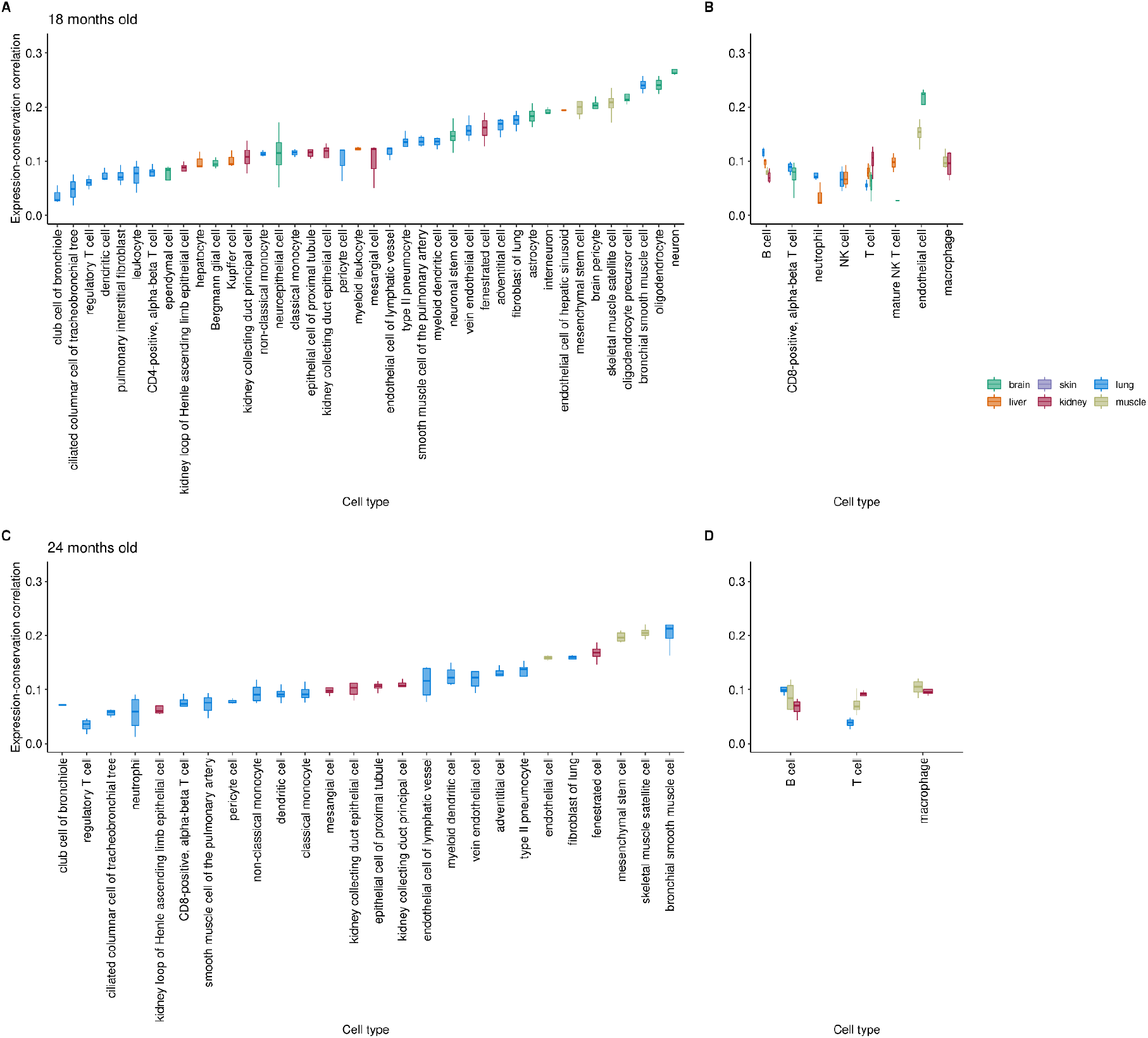
Variability of expression-conservation correlation levels across cell-type transcriptomes of 18-month-old (**A, B**) and 24-months-old (**C, D**) individuals. Each data point within bar plots represents Spearman’s correlation values for an individual, calculated using gene expression values averaged across cells belonging to that individual. Panels A, C and B, D show cell-types only sampled in one tissue and cell-types sampled from multiple tissues, respectively. Color-coding indicates the tissues which the cell-types were sampled from (see key at the top of the figure). Cell types with less than three correlation values (i.e. individuals) are excluded.

**Figure S5.**
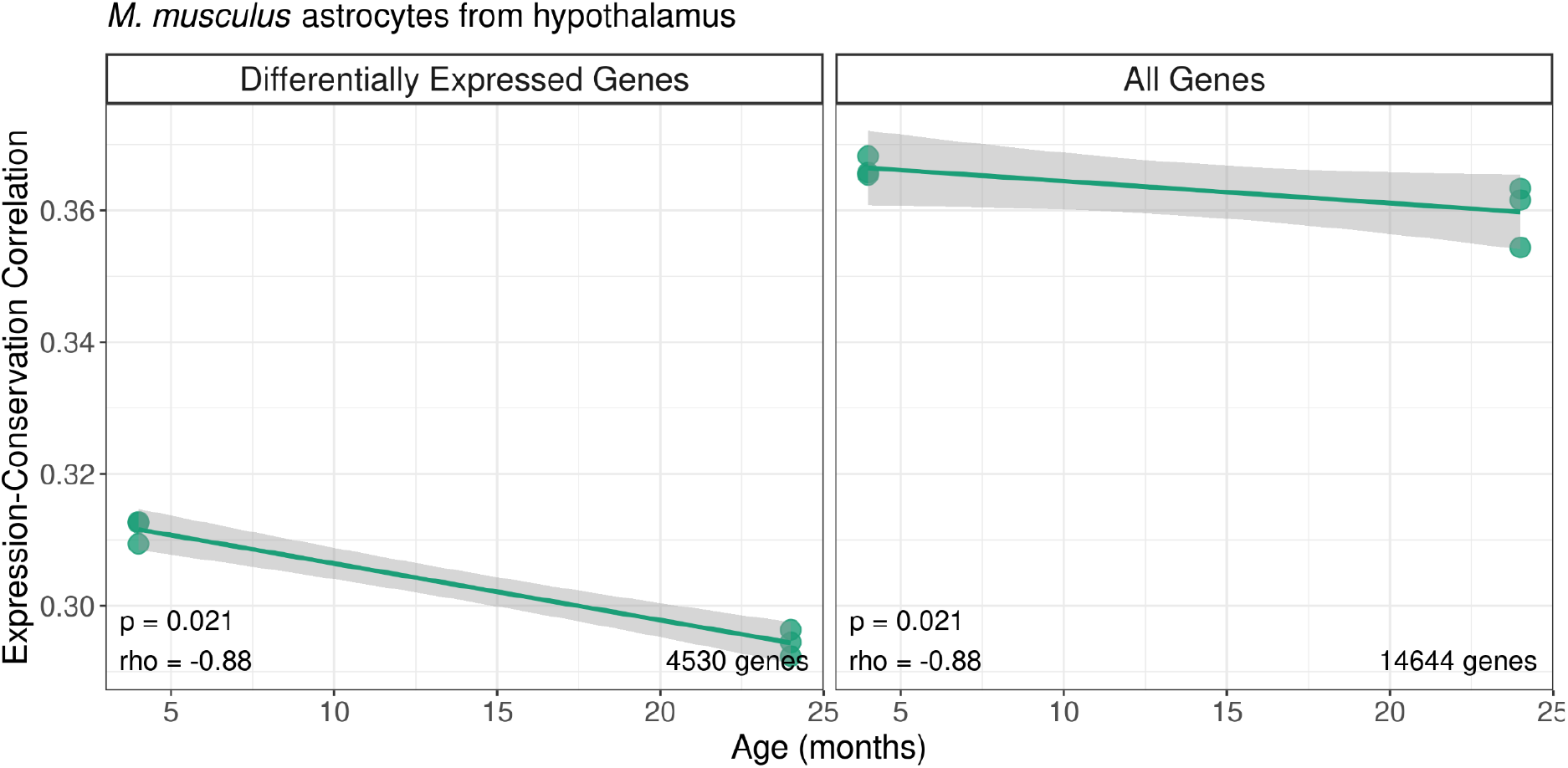
Age-related changes in expression-conservation correlation in *M. musculus* astrocyte transcriptome from the hypothalamus. The y-axis shows the Spearman correlation coefficient between expression level and protein sequence conservation metric across genes for each individual in this dataset (*n* = 6). The x-axis shows individual age. The left panel shows the analysis results using genes differentially expressed with age (*n* = 4,530), and the right panel shows the analysis results using the whole transcriptome (*n* = 14,644). *rho* and p-values in the inset indicate the results of the Spearman correlation between the expression-conservation correlation and age.

